# Millipede genomes reveal unique adaptation of genes and microRNAs during myriapod evolution

**DOI:** 10.1101/2020.01.09.900019

**Authors:** Zhe Qu, Wenyan Nong, Wai Lok So, Tom Barton-Owen, Yiqian Li, Chade Li, Thomas C.N. Leung, Tobias Baril, Annette Y.P. Wong, Thomas Swale, Ting-Fung Chan, Alexander Hayward, Sai-Ming Ngai, Jerome H.L. Hui

**Author notes:** = contributed equally (Zhe Qu, Wenyan Nong, Wai Lok So, Tom Barton-Owen, Yiqian Li). = corresponding author.

## Abstract

The Myriapoda including millipedes and centipedes is of major importance in terrestrial ecology and nutrient recycling. Here, we sequenced and assembled two chromosomal-scale genomes of millipedes *Helicorthomorpha holstii* (182 Mb, N50 18.11 Mb mainly on 8 pseudomolecules) and *Trigoniulus corallinus* (449 Mb, N50 26.78 Mb mainly on 15 pseudomolecules). Unique defense systems, genomic features, and patterns of gene regulation in millipedes, not observed in other arthropods, are revealed. Millipedes possesses a unique ozadene defensive gland unlike the venomous forcipules in centipedes. Sets of genes associated with anti-microbial activity are identified with proteomics, suggesting that the ozadene gland is not primarily an antipredator adaptation (at least in *T. corallinus*). Macro-synteny analyses revealed highly conserved genomic blocks between centipede and the two millipedes. Tight Hox and the first loose ecdysozoan ParaHox homeobox clusters are identified, and a myriapod-specific genomic rearrangement including Hox3 is also observed. The Argonaute proteins for loading small RNAs are duplicated in both millipedes, but unlike insects, an argonaute duplicate has become a pseudogene. Evidence of post-transcriptional modification in small RNAs, including species-specific microRNA arm switching that provide differential gene regulation is also obtained. Millipede genomes reveal a series of unique genomic adaptations and microRNA regulation mechanisms have occurred in this major lineage of arthropod diversity. Collectively, the two millipede genomes shed new light on this fascinating but poorly understood branch of life, with a highly unusual body plan and novel adaptations to their environment.

## Introduction

Arthropoda comprises the myriapods (millipedes and centipedes), crustaceans (shrimps, crabs, and lobsters), chelicerates (spiders, scorpions, and horseshoe crab), and insects. Collectively, these taxa account for the majority of described terrestrial and aquatic animal species (Figure 1A). While crustaceans, chelicerates, and insects have been the focus of intense research, the myriapods are comparatively less well studied, despite their great diversity and important ecological roles. In particular, arthropod genomic and transcriptomic information is highly uneven, with a heavy bias towards the crustaceans, chelicerates, and insects (Pisani et al 2013; Richards 2019). Yet, myriapods display many interesting biological characteristics, including a multi-segmented trunk supported by an unusually large number of legs. Centipede is Latin for ‘100 feet’, but centipedes actually have between 30 and 354 legs and no species has exactly 100 legs (Arthur and Chipman 2005). In contrast, millipede is Latin for ‘1000 feet’, and while millipedes include the ‘leggiest’ animal on Earth, no species has as many as 1000 legs, with the true number varying between 22 and 750 (Marek et al 2012). Myriapods were among the first arthropods to invade the land from the sea, during an independent terrestrialisation from early arachnids and insects, which occurred during the Silurian period ~400 million years ago (Minelli 2015). Today, the Myriapoda consists of ~16,000 species, all of which are terrestrial (Kenning et al 2017). Currently, just two myriapod genomes are available: the centipede *Strigamia maritima* (Chipman et al 2014), and a draft genome of the millipede *Trigoniulus corallinus* (Kenny et al 2015). Consequently, the myriapods, and particularly the millipedes, present an excellent opportunity to improve understanding of arthropod evolution and genomics.

**Figure 1.**
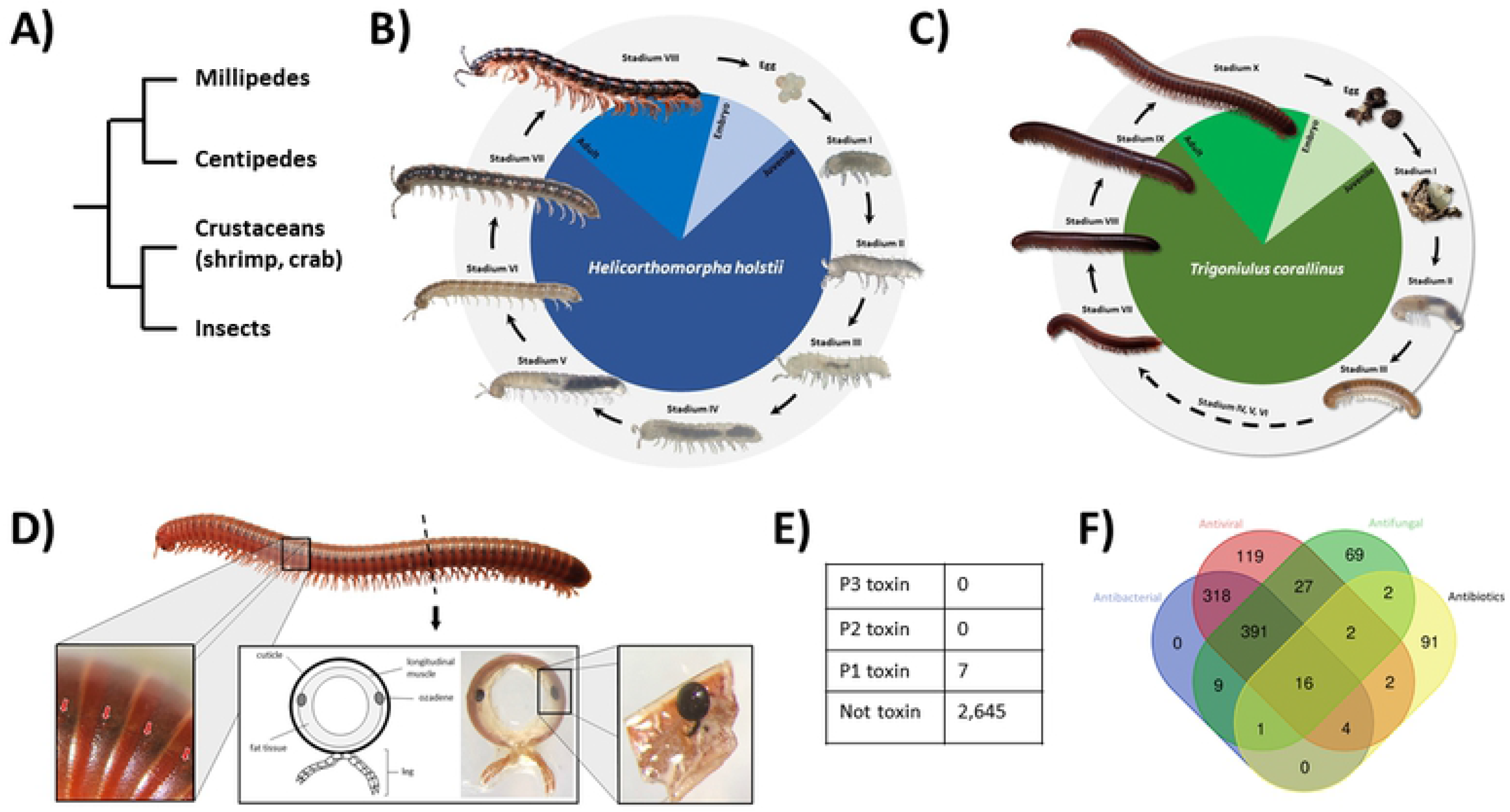
A) Schematic diagram showing the phylogeny of myriapods, crustaceans, and insects; B) Life cycle of polydesmid millipede *Helicorthomorpha holstii*; C) Life cycle of rusty millipede *Trigoniulus corallinus*; D-F) Ozadene defensive gland of millipede *T. corallinus* and its proteomic analyses.

Millipedes compose the class Diplopoda, a highly diverse group containing more than 12,000 described species, and the third largest class of terrestrial arthropods after insects and arachnids (Adis 2002). Millipedes are highly important components of terrestrial ecosystems, especially with reference to their roles in the breakdown of organic plant materials and nutrient recycling. In contrast to centipedes which have one pair of legs per body segment, individual body segments are fused in pairs in millipedes, resulting in a series of double-legged segments. The typical millipede body plan consists of the head, collum, and trunk (with varying numbers of diplo-segments). The primary defense mechanism of millipedes is to curl into a coil, while a unique secondary defense system in some species involves emitting toxic liquids or gases from the ozadene gland, via ozopores located on each side of the metazonite (the posterior portion of diplosegment)(Enghoff 1993).

The polydesmid millipede *Helicorthomorpha holstii* (Polydesmida) and the rusty millipede *Trigoniulus corallinus* (Spirobolida) were chosen in this study to represent two major lineages from the 16 millipede orders. Both species originate in Asia and are now cosmopolitan species. *H. holstii* undergoes development with fixed numbers of legs and segments that increase in every stadium after each molt, and will complete seven juvenile stadia before reaching sexual maturity at stadium VIII (adult) (Figure 1B). Conversely, *T. corallinus* undergoes development with variable numbers of new segments and legs added in the initial molts, with no further segments developing after reaching stadium X (adult)(Figure 1C).

Here we present two high-quality *de novo* reference genomes close to the chromosomal-assembly level, for the Asian polydesmid millipede *H. holstii* and the spirobolid rusty millipede *T. corallinus* (Table 1). With reference to these genomes, we reveal the basis of a unique defence system, genomic features, and gene regulation in millipedes, not observed in other arthropods. The genomic resources we develop expand the known gene repertoire of myriapods and provide a genetic toolkit for further understanding of their unique adaptations and evolutionary pathways.

**Table 1.**
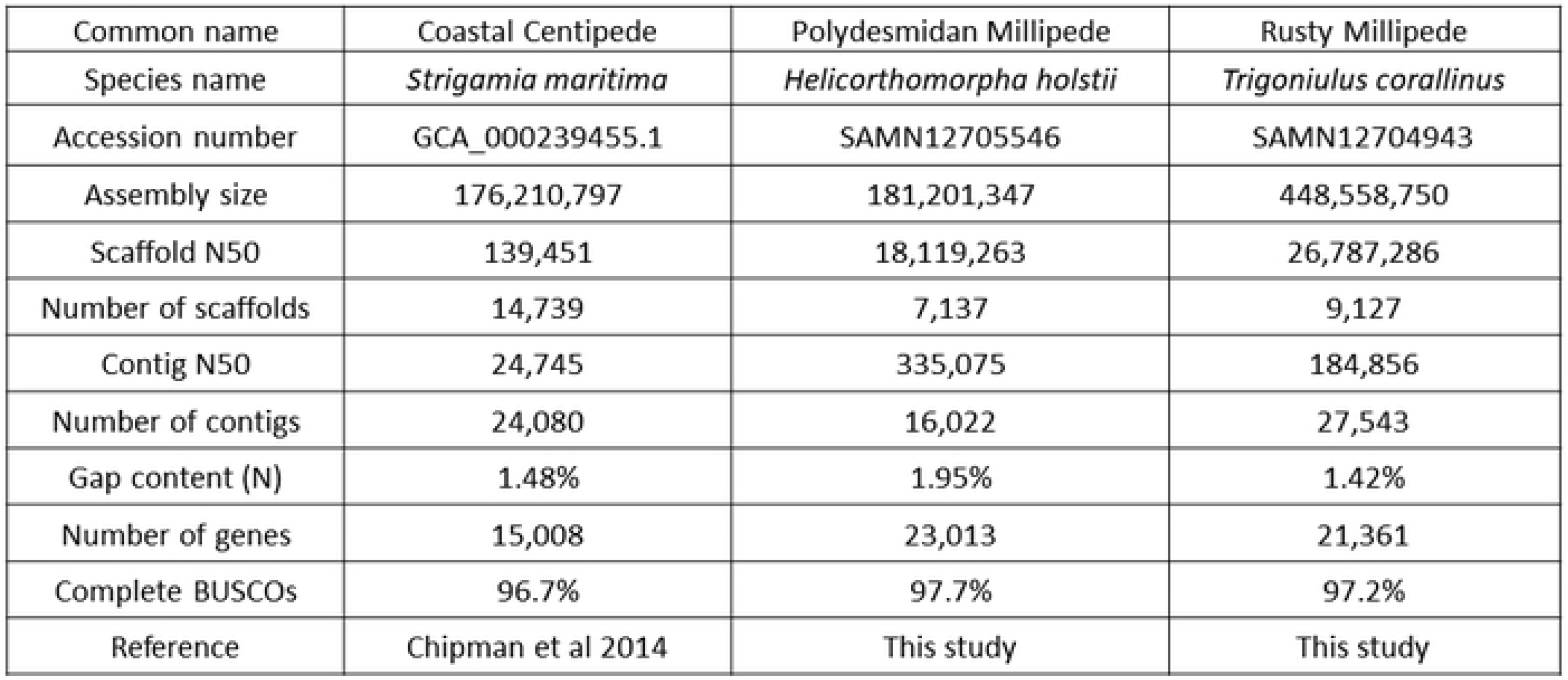
Comparison of myriapod genome assembly quality.

## Results and Discussion

### High quality genomes of two millipedes

Genomic DNA was extracted from single individuals of two species of millipedes, including polydesmid millipede *H. holstii* (Figure 1B) and the rusty millipede *T. corallinus* (Figure 1C), and sequenced using the Illumina short-read and 10X Genomics linked-read sequencing platforms (Supplementary information S1, Table 1.1.1-1.1.2). Hi-C libraries were also constructed for both species and sequenced on the Illumina platform (Supplementary information S1, Figure S1.1.1-1.1.2). Both genomes were first assembled using short-reads, followed by scaffolding with Hi-C data. The *H. holstii* genome assembly is 182 Mb with a scaffold N50 of 18.11 Mb (Table 1). This high physical contiguity is matched by high completeness, with a 97.2 % complete BUSCO score for eukaryotic genes (Table 1). The *T. corallinus* genome is 449 Mb with a scaffold N50 of 26.7 Mb, 96.7 % BUSCO completeness, (Table 1). 23,013 and 21,361 gene models were predicted for the *H. holstii* and *T. corallinus* genome assemblies, respectively (Table 1). Majority of the sequences assembled for the *H. holstii* and *T. corallinus* genomes are contained on 8 and 17 pseudomolecules respectively (Supplementary information S1, Figure S1.1.1-1.1.2), representing the first close to chromosomal-level genomes for myriapods.

### Ozadene defensive gland

Many millipedes possess ozadenes, specialised glands that contain chemicals such as alkaloids, quinones, phenols, or cyanogenic compounds used in defence against predators. The structure of the julid-type gland in *T. corallinus* consists of sacs bordered by secretory cells lined with cuticle, and an efferent duct opening laterally on the body surface via a small ozopore (Figure 1D). Discharge of defensive compounds is accomplished by contraction of valvular muscle and simultaneous compression of the sac. 2,652 peptides were identified in the *T. corallinus* ozadene gland by mass spectrometry. Despite a vast number of peptides being identified, only 7 of them were predicted as P1 toxins (i.e. with low confidence, Figure 1E, Supplementary information S1, Figure S1.3.1, Table 1.3.3, Supplementary information S3). These data suggest that the millipede ozadene gland (at least for *T. corallinus*), is not adapted to produce toxins, unlike the venomous forcipules present in centipedes.

The question then becomes, what is the function of the millipede ozadene gland? Gene ontology and KEGG pathway analyses of the remaining 2,645 non-toxin peptides were performed (Supplementary information S1, Table 1.3.4, and S4), and identified a total of 1,051 proteins involved in antibacterial, antifungal, and antiviral biosynthesis (Figure 1F, Supplementary information S1, Figure S1.3.2, and S4). These data and analyses suggest one of the main functions of millipede ozadene (at least in *T. corallinus*), is to provide defence against pathogenic microorganisms rather than being primarily an antipredator adaptation.

### Conserved synteny between myriapod genomes

A major reason for the broad significance of millipede genomic resources is that myriapods serve as the outgroup to the Insecta, which is the largest group of described animal species. Thus, comparisons between millipedes and insects allow us to address a major outstanding question in animal evolution, specifically, how differential regulation of gene function facilitated the evolution of greatly divergent body plan morphologies.

Conservation of large-scale gene linkage has been previously detected between the centipede *Strigamia maritima* and the amphioxus *Branchiostoma floridae* at a higher level than with any insect, proving evidence that the last common ancestor of arthropods retained significant synteny with the last common ancestor of bilaterians (Chipman et al 2014). To understand the genomic rearrangement patterns among and between the diplopods and chilopods, conserved synteny analyses between the two millipedes and the centipede *S. maritima* were performed. As expected, higher conserved synteny blocks can be detected between the two millipedes than between millipede and the centipede (Figure 2). A higher level of large-scale gene linkage is also observed between *T. corallinus* and *S. strigamia* than between *H. holstii* and *S. maritima* (Figure 2). To further shed light on the situation, we have also compared the syntenic relationships between the three myriapod genomes to that of the human, and found that both millipede genomes share more syntenic blocks with human than human genome with the centipede genome (Figure 2).

**Figure 2.**
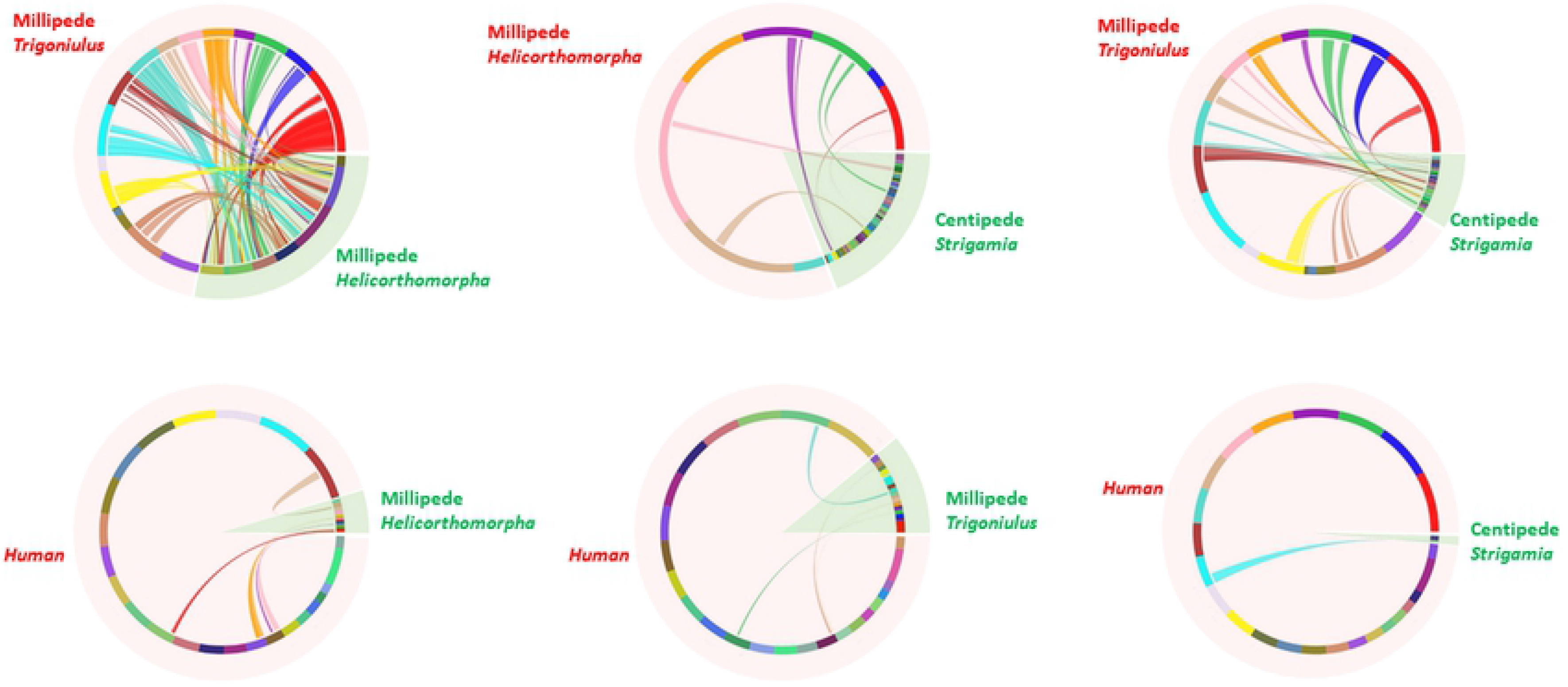
Synteny comparisons of myriapod and human genomes.

### Millipede homeobox gene and cluster

Homeobox genes are an ideal candidate to study body plan evolution, as they are conserved gene expression regulators in animals. We first systematically compared the homeobox gene content of all available insect genomes to the three myriapod genomes (Supplementary information S1, Figure S1.3.4). Given the varying genome quality of the insects being compared, we adopted a conservative approach by only confidently scoring gene gains, rather than gene losses, using the rationale of revealing the presence of orthologous homeobox genes in closely related lineages. The three myriapod genomes have undergone 3 lineage-specific duplications of common homeobox genes (Otx, Barhl, Irx), suggesting their suitability for making a comparison to the insects (Supplementary information S1, Figure S1.3.4).

Hox gene clusters are renowned for their role in the developmental patterning of the anteroposterior axis of animals. In both *H. holstii* and *T. corallinus* genomes, intact Hox clusters containing orthologues of most arthropod Hox genes are recovered except Hox3 gene, and are expressed in early developmental stages (Figure 3, Supplementary information S1, Figure S1.3.3, S1.3.6, S1.3.7). In the *H. holstii* genome, no Hox3 orthologues could be identified, and two Hox3 genes were located together on a different scaffold to the Hox cluster scaffold in *T. corallinus*. This situation mirrors that in the centipede *S. maritima* (Chipman et al 2014). Based on our phylogenetic analyses (Supplementary information S5 and S6), we suggest that the genomic “relaxation” of Hox3 from the intact tight Hox clusters may have occurred in the ancestor of all myriapods (Figure 3).

**Figure 3.**
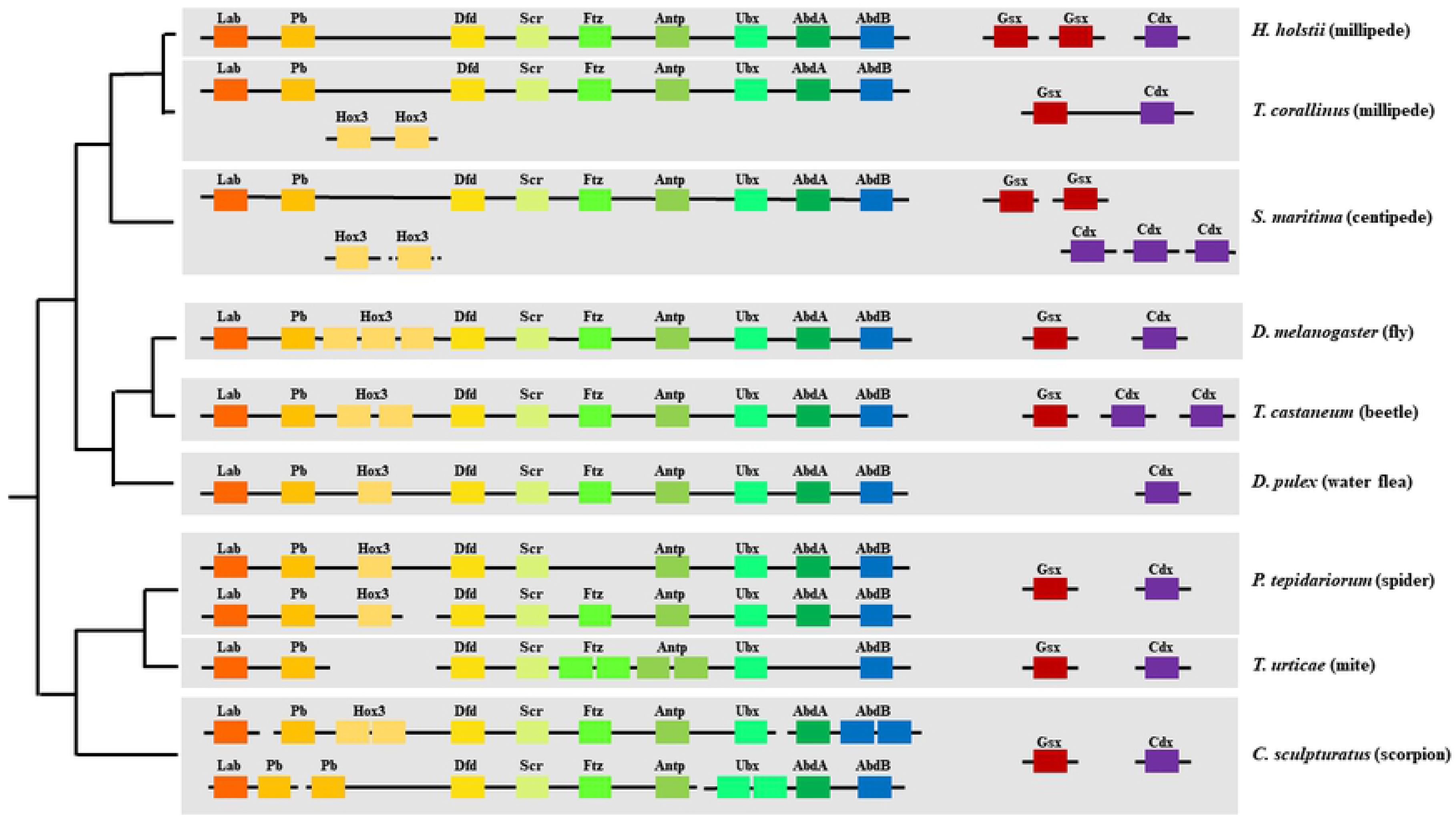
Hox and ParaHox gene cluster genomic organisations in millipedes and other arthropods.

Similar to the situation in *S. maritima*, Eve orthologue is closely linked to the Hox clusters in both millipede genomes. In addition, we have also discovered the linkage of other ANTP-class homeobox gene members to the Hox-Eve in the millipede genomes, including Abox, Exex, Dll, Nedx, En, Unpg, Ro, Btn (Supplementary information S1, Figure S1.3.5, S1.3.8). Whether this represents a difference in genome quality between the two millipede genomes (N50 = 26.7 Mb and 18.1 Mb) and the centipede genome (Chipman et al 2014, N50 = 139kb), or a true genomic content difference between these myriapod lineages, remains to be tested through improvements to centipede genomic resources.

The ParaHox cluster is the paralogous sister of the Hox cluster, and contains an array of three ANTP-class homeobox genes: Gsx/ind, Xlox/Pdx, and Cdx/cad together patterning the brain and endoderm formation in bilaterians (Brooke et al 1998). In general, the genomic linkage of ParaHox cluster genes has been lost in all investigated ecdysozoans (Hui et al 2009). A loosely linked ParaHox cluster of Gsx and Cdx is found in the millipede *T. corallinus*, representing the first ecdysozoan ParaHox cluster (Figure 3). The ParaHox genes are expressed mainly during early development, and Gsx is also expressed during late developmental stage (Supplementary information S1, Figure S1.3.7). Given ParaHox clustering has been identified in the lophotrochozoans and deuterostomes (Brooke et al 1998; Hui et al 2009), our data here provide evidence that the arthropod and ecdysozoan ancestors contained clustering of ParaHox genes rather than having disintegrated ParaHox cluster as previously thought.

Other homeobox gene clusters are also identified and compared, including the NK cluster and the Irx cluster (Supplementary information S1, Figure S1.3.6, S1.3.8). Collectively, these examples highlight the importance of the novel genomic resources presented here, to: 1) reconstruct the arthropod ancestral situation, providing different interpretations given lineage-specific modifications, and, 2) understand the functional constraints in extant lineages, such as the relaxation in Hox3 and Xlox in the ecdysozoan Hox and ParaHox clusters.

### Transposable elements

Transposable elements (TEs) are almost ubiquitous components of eukaryotic genomes, often accounting for a large proportion of an organism’s genome (Chénais et al 2012). Among the metazoans, the phylum Arthropoda is a particular focus for TE research. However, Myriapoda are the only major branch of Arthropoda for which knowledge of TEs remains extremely poor. Here, we examined the repeat content of one centipede and two millipede genomes to perform the first comparative investigation of TEs in the Myriapoda. As for other major arthropod groups (Petersen et al 2019), we find considerable variation in the total genomic contribution and composition of TEs among myriapod genomes.

TEs comprise 19-47% of assembled genomic content among the three available myriapod genomes, with total repeat content varying between 19-55% (Figure 4: Repeat Content; Supplementary information S1, Table 1.3.2). In the spirobolid millipede *T. corallinus*, repeats account for more than half of the assembled genome (55%, Supplementary information S2), which is of interest since the genome of *T. corallinus* is more than double the length of either of the other two myriapod genomes (Figure 4: Repeat Content), suggesting that TEs have played a role in genome size evolution in this species. In contrast, repeat content is 40% in the geophilomorph centipede *S. maritima*, and just 19% in the polydesmid millipede *H. holstii*, demonstrating considerable variation among lineages (55%, Supplementary information S2). It is unclear what mechanisms are responsible for generating this variation, however, similar levels of variability in TE content are typical among species in other invertebrate genomes.

**Figure 4.**
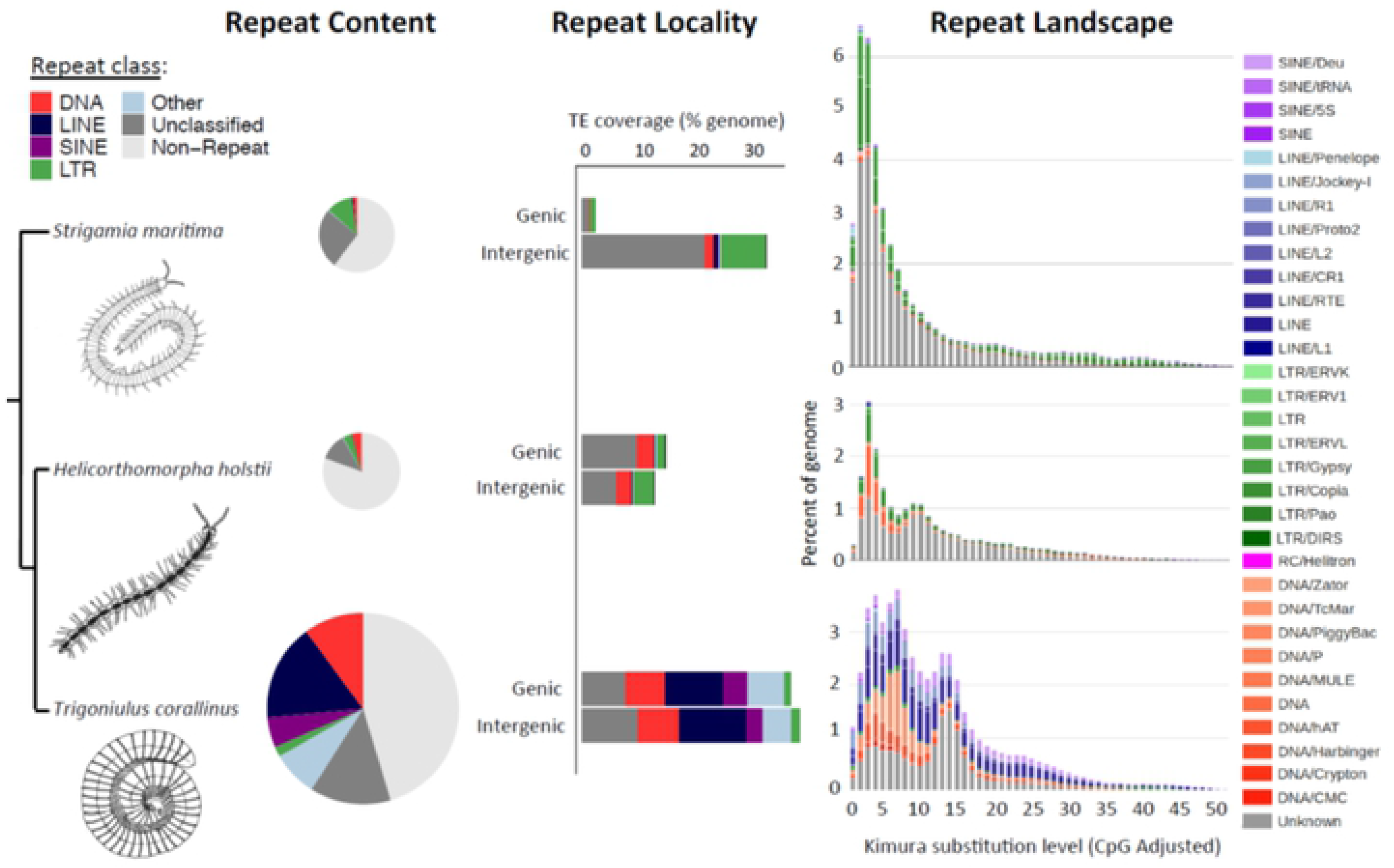
Transposable element content, genomic locality, and estimates of accumulation history for sequenced members of the Myriapoda. Phylogenetic relationships among taxa are indicated on the left-hand side of the figure, alongside schematics of each myriapod species. From left to right: (i) Pie charts in proportion to assembled genome size illustrating the relative contribution to myriapod genomes from each major repeat class; (ii) Stacked bar charts illustrating the proportion of each repeat class found in genic (≤2kb from an annotated gene) versus intergenic regions (>2kb from an annotated gene) for each myriapod species, expressed as a percentage of the total assembled genome; (iii) Repeat landscape plots illustrating transposable element accumulation history for each myriapod genome, based on Kimura distance-based copy divergence analyses, with sequence divergence (CpG adjusted Kimura substitution level) illustrated on the x-axis, percentage of the genome represented by each transposable element type on the y-axis, and transposon type indicated by the colour chart on the right-hand side.

The main repeat types identified differ considerably among the available myriapod genomes. Complex retrotransposons, particularly *copia*-like and *gypsy*-like long-terminal repeat (LTR) TEs are the dominant TE class present in the genome of the centipede *S. maritima*, while Maverick DNA TEs are also dominant in the genome of the millipede *H. holstii* (Supplementary information S2). Meanwhile, a more cosmopolitan set of TEs are identified in the genome of *T. corallinus*, including SINEs, LINEs, DNA TEs and *gypsy-like* LTR TEs (Supplementary information S2).

TE sequences are distributed evenly across genic and intergenic regions in both millipede genomes (Figure 4: Repeat Locality). The frequent annotation of TEs in close proximity to genes raises the possibility that TEs may have played a particular role in host evolution in millipedes, since TEs are well known contributors of genomic novelty to host genomes (e.g. Feschotte 2008; Kidwell and Lisch 2000; Schrader and Schmitz 2019). Strikingly, in comparison TEs are largely excluded from genic neighbourhoods in the centipede genome (Figure 4: Repeat Locality). These patterns suggest that millipedes may be a particularly furtile group for future studies on host-TE interactions.

Analyses of sequence divergence among annotated TEs suggest that all three myriapod genomes have experienced recent spikes in TE activity, however, the specific pattern of activity difference among species (Figure 4: Repeat Landscapes). Strikingly, there is evidence of a particularly large and recent expansion of LTR TEs in the centipede *S. maritima*, but very limited evidence for activity prior to this, suggesting a recent invasion of the genome by copia and gypsy LTR TEs (Figure 4: Repeat Landscapes). In the millipede *H. holstii*, there is evidence of a much more modest recent expansion of both LTR TEs and DNA TEs, while there is evidence of a more prolonged expansion including SINEs, LINEs, and DNA TEs, but very little LTR TE activity, in the millipede *T. corallinus* (Figure 4: Repeat Landscapes).

Taken together, our findings suggest that TEs have played a significant role in the shaping of myriapod genomes, implying that myriapods represent a rich group for future studies on host-TE interactions.

### Specific duplication of argonaute protein

Another ideal candidate to understand how animals evolve are the small RNAs and their associated machineries. Small RNAs are also conserved gene expression regulators in animals, and their study will reveal hidden layers of gene regulation. For instance, the well-known mature microRNAs are 21-23 nucleotide non-coding RNAs that regulate gene expression and translation, usually by binding onto the 3’ UTRs of target mRNAs to achieve post-transcriptional inhibition, either by suppressing translation or inducing mRNA degradation (Cao et al 2017; Qu et al 2018; Figure 5A). Despite the finding that the biogenesis pathways of microRNAs and other small RNAs are relatively conserved in animals, modifications of the small RNA machineries have been found to alter small RNA regulation and thus contribute to rewiring of genetic networks. For instance, the placozoan *Trichoplax adhaerens* has lost Piwi, Pasha, and Hen1 genes from its genome, where no microRNA is found to be produced (Grimson et al 2008).

**Figure 5.**
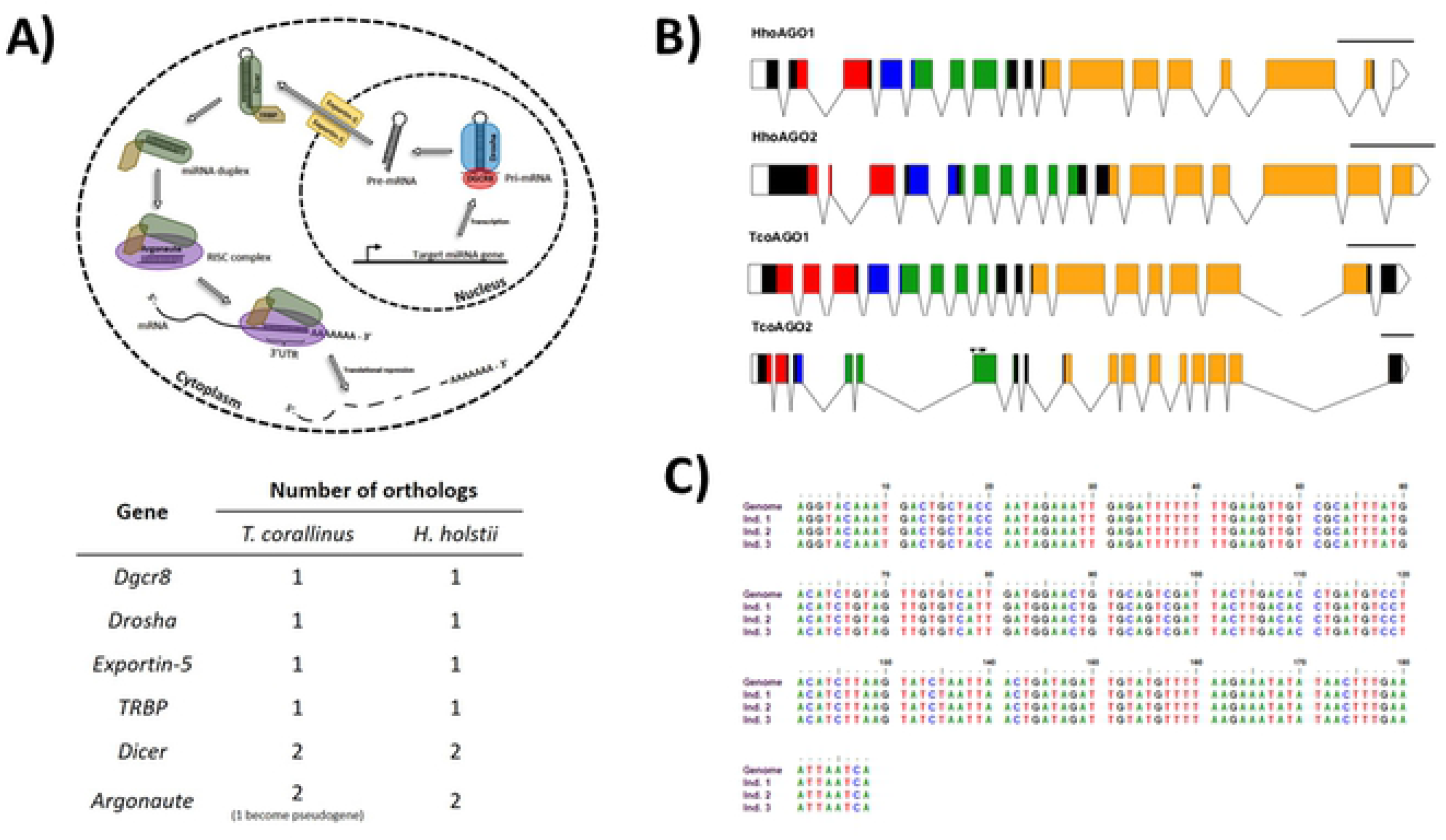
A) Schematic diagram showing the biogenesis pathway of microRNAs (upper) and table summarising the number of gene copies contained in each millipede genome (lower); B) Schematic diagram showing the duplicates of Argonaute (Ago) gene in the two genomes. Conserved domains of AGO - ArgoN (red), ArgoL (blue), PAZ (green) and PIWI (orange). Inverted triangles, in TcoAGO2, indicate the position of multiple stop codons found in the corresponding gene sequence. Scale bar = 500 nucleotides; C) Confirmation of TcoAGO2 pseudogene. PCR and Sanger sequencing were carried out on genomic DNA collected from three *Trigoniulus corallinus* individuals that are not used for genome sequencing.

In the two millipede genomes generated here, all genes responsible for small RNA machinery were identified, along with an unusual duplication of the Argonaute (Ago) gene, while the other biogenesis components remain the same (Figure 5A, supplementary information S1, Figure S1.3.9-1.3.11). In insects, it is well known that there are also two Ago forms, and for instance, in the fly *Drosophila melanogaster*, the dominant arm of the precursor microRNA can be sorted into Ago1 to direct translational repression. Meanwhile, the other arm as well as small-interfering RNA (siRNA) can be sorted into Ago 2 to direct transcriptional degradation (Czech et al 2009; Ghildiyal et al 2009; Okamura et al 2008, 2009; Yang et al 2011). Nevertheless, phylogenetic analyses suggested the two Ago forms in the two millipede genomes are lineage-specific, and did not share the duplication event that occurred in insects (Supplementary information S1, Figure S1.3.9). In addition, one Ago in *T. corallinus* (which we named Ago2) has become a pseudogene (Figure 5B). To test that this is not due to genome assembly error, nor due to single individual mutation, we have also carried out PCR and Sanger sequencing on three other individuals, and confirmed this mutation (Figure 5C). Whether the duplicated Ago are functional in millipedes remains unclear. However, this unusual duplication of the small RNA machinery in millipedes reveals that the situation in insects was secondarily evolved, rather than shared with the duplication that occurred in the arthropod ancestor.

### MicroRNA regulate homeobox genes and arm switching

To understand how posttranscriptional regulators have evolved in this special lineage, small RNA transcriptomes were obtained from the eggs, juveniles, and adults of *H. holstii* and *T. corallinus* (Supplementary information S1, Table 1.1.3-1.1.4). Using stringent criteria to annotate microRNAs which were supported by small RNA reads, a total of 59 and 58 conserved microRNAs were identified in the genomes of *H. holstii* and *T. corallinus* respectively (Supplementary information S7-S9). This number is comparable to the 58 microRNAs identified in the centipede *S. maritima* (Chipman et al 2014). In addition to the conserved microRNAs, 43 and 10 novel lineage-specific microRNAs could further be identified in millipedes *H. holstii* and *T. corallinus* respectively, and only one of them is conserved between the two millipedes (Supplementary information S1 Figure S1.3.13, Supplementary information S7-S9). Whether these novel microRNAs have contributed to the unique adaptation of millipedes deserves further explorations.

In the centipede *S. maritima*, a homologue of miR-125, which is a member of the ancient bilaterian miR-100/let-7/miR-125 cluster, could not be identified (Chipman et al 2014; Griffiths-Jones et al 2011). However, miR-125, could be identified in both millipede genomes, suggesting a lineage-specific loss in the centipede (Figure 6A). In addition, our two high-quality millipede genomes allowed us to reveal conserved microRNA clusters, including miR-100-let-7-125, miR-263-96, miR-283-12, miR-275-305, miR-317-277-34, miR-71-13-2, miR-750-1175 and miR-993-10-iab4/8, as in most other arthropods (Supplementary information S9). Previously, miR-283 has been identified in pancrustaceans only, but it could be identified in the two millipede genomes presented here (Supplementary information S9). Moreover, miR-96 and miR-2001 could be identified in the two millipedes, but not in *S. maritima* (Supplementary information S9). These examples highlight the importance of having multiple high-quality myriapod genomes for comparison to properly understand the evolution of post-transcriptional regulators.

**Figure 6.**
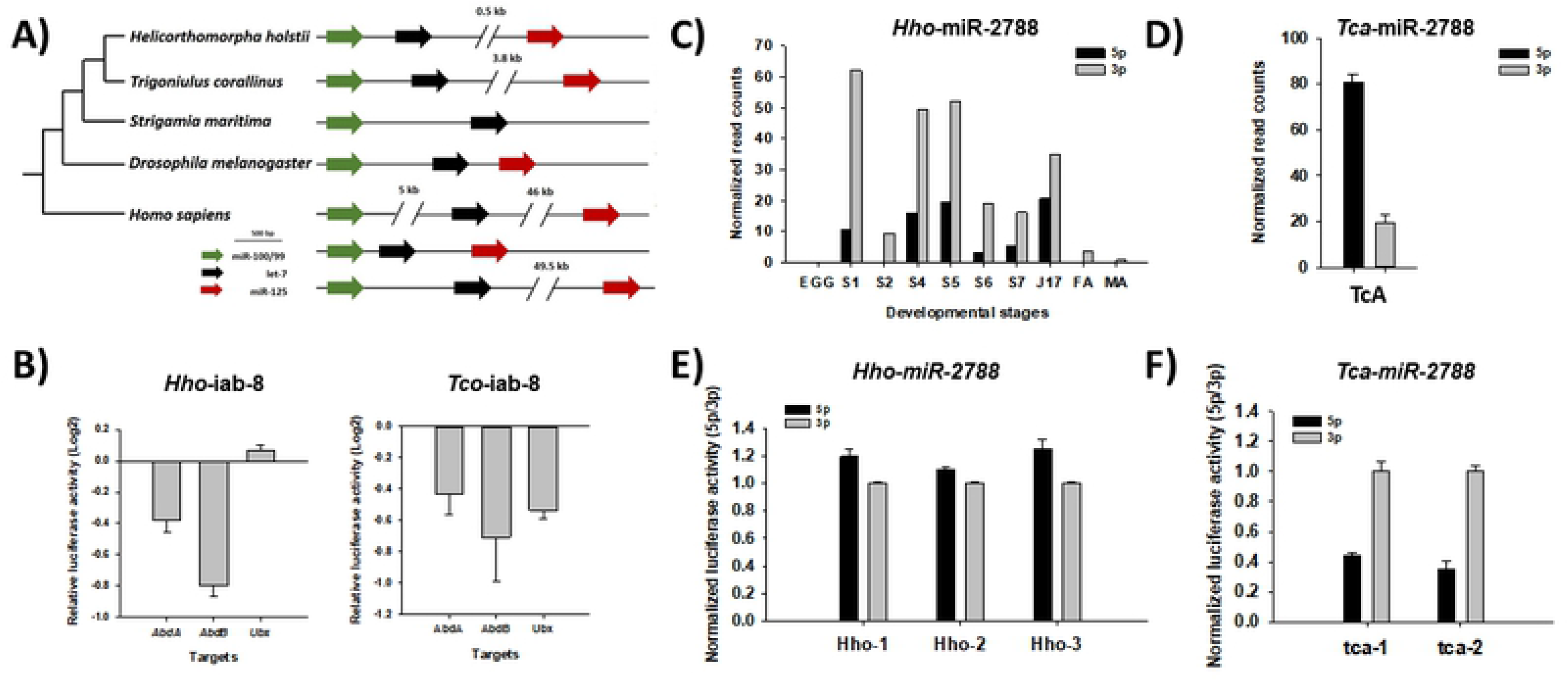
A) Genomic organisation of miR-100/let-7/miR-125 clusters in various animals; B) Luciferase assays showing the repression activities of Hox genes by miR-iab-8 in both millipedes; D-E) Small RNA read counts of miR-2788 in different developmental stages in millipede *Helicorthomorpha holstii* and in TcA cell line of beetle *Tribolium castaneum;* Abbreviations: S1-S7: Stadium 1-7, J17: Juvenile, FA: adult female, MA: adult male, TcA: TcA cell line; I-J) Luciferase activity showing the differential arm target (i.e. miR-2788-5p and −3p sensor) repression ability between miR-2788 carrying different flanking sequence of *H. holstii* and *T. castaneum*.

We further explored how conserved microRNAs may modulate gene regulatory networks among arthropod lineages. In insects, the bidirectionally transcribed microRNA iab-4/iab-8 locus is renowned for its regulation of the functions of its flanking Hox genes in the genomic cluster (Hui et al 2013). In both millipedes *H. holstii* and *T. corallinus*, the microRNA iab-4/iab-8 locus is located between Hox genes *abd-A* and *abd-B* similar to the situation in insects. In the cell-based dual-luciferase reporter assay to test Hox gene targets targeted by iab-8 in the two millipedes, we found that the posterior Hox genes can be downregulated in the two millipedes (abd-A and abd-B by *H. holstii* iab-8, abd-A, abd-B and Ubx by *T. corallinus* iab-8) as in the situations in insects (Figure 6B). These data further established the regulation of Hox genes Ubx and abd-A by intergenic microRNA iab-8 in the most recent common ancestor of insects and myriapods.

Many miRNA loci produce significant quantities of mature miRNAs from both arms with different amounts (5p or 3p), and RISC-loaded miRNA will then bind complementary to the 3’UTR of mRNA. These result in the suppression of gene expression by either, promoting mRNA cleavage, translational repression, or decay due to deadenylation (Ghildiyal and Zamore, 2009). In animals, complementary base pairing of nucleotides 2 to 8 in the 5’ of the miRNA (also known as the seed region) is pivotal and efficient for targeting to mRNA (Krol et al 2010; Griffiths-Jones et al 2011). Since the sequences of alternative mature miRNAs derived from opposite complimentary arms are different, mature miRNAs derived from the same hairpin will also regulate distinct sets of genes (Marco et al 2010, 2012; Berezikov 2011; Griffiths-Jones et al 2011). Interestingly, the choice of dominant arm expression can be swapped at different situations, a termed known as microRNA arm switching, such as a-b) developmental stages, tissues (e.g. Ro et al 2007; Ruby et al 2007; Glazov et al 2008; Chiang et al 2010; Jagadeeswaran et al 2010; Cloonan et al 2011; Biryukova et al 2014; Gong et al 2014; Pundhir and Goodkin 2015); c) pathological statuses (e.g. gastric cancer, Li et al 2012); and d) species (e.g. de Wit et al 2009; Marco et al 2010; Griffiths-Jones et al 2011; Brawand et al 2014; Sadd et al 2015). Comparing conserved microRNAs between the two millipedes and also all available insect genomes with small RNA sequencing data, we found multiple cases of microRNAs undergoing microRNA arm switching, including let-7 and miR-277 (Supplementary information S1 Figure S1.3.13, S1.3.15, Supplementary information S9).

To decipher how microRNA arm switching could potentially contribute to evolution between myriapods and insects, we focused on the two microRNAs iab-8 and miR-2788, which were previously only known from insects but are now also known to be conserved in centipede and millipedes (Chipman et al 2014; this study). Using the sensor assays, we found no obvious arm target repression preference of iab-8 of the two millipedes, but the *Drosophila* iab-8 have higher 5p dominant arm and target repression ability (Supplementary information S1 Figure S1.3.14). These data suggest that different arm usage of iab-8 has evolved between insects and myriapods. In the small RNA sequencing of the beetle *Tribolium castaneum* cell line and different developmental stages of millipedes *H. holstii*, miR-2788 shows different arm preferences (5p dominance in *T. castaneum* and 3p dominance in *H. holstii*) (Figure C-D). To test the targeting properties of *T. castaneum* and *H. holstii* miR-2788, a set of luciferase reporters containing perfect target sites for both 5p and 3p mature miR-2788 were constructed and co-transfected into S2 cells, with expression constructs driving production of either species’ miR-2788. Repression of target sites for the 5p and 3p arms of each hairpin was found to correlate with their relative production as determined by next-generation sequencing (Figure 6E–F). As sequence conservation outside microRNA hairpin sequences in the flanking sequences have been identified across species (Kenny et al 2015), and the dominant usage of arms of microRNA candidates such as miR-10 in insects are not governed by thermodynamics (Griffiths-Jones et al 2011), microRNA flanking sequence has been suggested as a potential candidate to govern arm switching (Griffiths-Jones et al 2011; Kenny et al 2015). The flanking sequences of both *T. castaneum* and *H. holstii* miR-2788 were deleted and transfected to S2 cells for luciferase reporter assays, and our results revealed that the same dominant arm is associated with various flanking sequences (Figure 6E-F). This suggests that the governance of microRNA arm switching could be under multiple mechanisms, and candidate-specific during evolution, presenting an additional means of adaptation.

## Conclusions

The two millipede chromosomal-level genomes provided in this study expand the gene repertoire of myriapods and arthropods. The phylogenetic position of myriapods within the arthropods provides a genetic toolkit for the reconstruction of evolutionary histories in insect and arthropod ancestors, as well as understanding their unique adaptations.

## Materials and Methods

### Genome and transcriptome sequencing and assembly

Genomic DNA was extracted from a single individual for each millipede, while mRNA and small RNA were extracted from a range of tissues. Different sequencing platforms were used to sequence genomic DNA, mRNA and small RNA. *De novo* genome and transcriptome assemblies were carried out following the methods described in the Supplementary information S1.

### Annotations and evolutionary analyses

Gene model predictions were carried out with the support of mRNA. The identities, genomic locations, and expression of different gene families and small RNAs were analysed. Details are provided in the Supplementary information S1.

## Acknowledgements

This work was supported by Hong Kong Research Grants Council (RGC) General Research Fund (GRF) (14103516, 14100919) and The Chinese University of Hong Kong (CUHK) Direct Grant (4053248) to JHLH. AH is supported by a Biotechnology and Biological Sciences Research Council (BBSRC) David Phillips Fellowship (BB/N020146/1). TB is supported by a studentship from the BBSRC-funded South West Biosciences Doctoral Training Partnership (BB/M009122/1). WLS, YL, CL are supported by studentships from The Chinese University of Hong Kong. The funders had no role in study design, data collection and analysis, decision to publish, or preparation of the manuscript. The authors would like to thank K. Chong, N. Kenny, C.Y. Lee, H.Y. Yip, and C. Yu for the initial works on millipede collections and/or materials extraction.

## Competing Interests

None of the authors have any competing interests.

## Availability of data and materials

The genomic and transcriptomic data generated in this study have been deposited to NCBI under BioProjects PRJNA564202 (*Helicorthomorpha holstii*) and PRJNA564195 (*Trigoniulus corallinus*).

## Supplementary Information

S1. Supplementary methodology and data.

S2. Transposable elements in the two millipede genomes.

S3. List of toxin-like proteins predicted in the myriapod genomes.

S4. List of proteins identified in the *Trigoniulus corallinus* ozadene gland.

S5. Homeobox gene sequences annotated in the two millipede genomes.

S6. Homeobox gene tree.

S7. *Helicorthomorpha holstii* microRNA structures.

S8. *Trigoniulus corallinus* microRNA structures.

S9. The microRNA contents and arm usage in *Tribolium castaneum* and *Helicorthomorpha holstii*.

